# The Mitochondrial DNA Control Region might have useful Diagnostic and Prognostic Biomarkers for Thyroid Tumors

**DOI:** 10.1101/291435

**Authors:** Rifat Bircan, Hülya Iliksu Gözü, Ulu Esra, Şükran Sarikaya, Aylin Ege Gül, Duygu Yaşar Şirin, Serhat Özçelik, Cenk Aral

**Affiliations:** Namık Kemal University, Arts and Sciences Faculty, Department of Molecular Biology & Genetics, Tekirdağ/TURKEY; Marmara University, School of Medicine, Department of Endocrinology and Metabolism, İstanbul/TURKEY; Kartal Lütfi Kırdar Education & Research Hospital, Department of Pathology, İstanbul/TURKEY; Hardarpaşa Education andResearch Hospital, Section of Endocrinology and Metabolism, Istanbul/TURKEY

**Keywords:** Mitochondrial DNA, Control region, D310, D514 CA repeat, T16189C, Thyroid Nodule, Papillary Thyroid Cancer

## Abstract

**Background:** It is currently present in the literature that mitochondrial DNA (mtDNA) defects are associated with a great number of diseases including cancers. The role of mitochondrial DNA (mtDNA) variations in the development of thyroid cancer is a highly controversial topic. In this study, we investigated the role of mt-DNA control region (CR) variations in thyroid tumor progression and the influence of mtDNA haplogroups on susceptibility to thyroid tumors.

**Material & method:** For this purpose, totally 108 hot thyroid nodules (HTNs), 95 cold thyroid nodules (CTNs), 48 papillary thyroid carcinoma (PTC) samples with their surrounding tissues and 104 healthy control subject’s blood samples were screened for entire mtDNA CR variations by using Sanger sequencing. The obtained DNA sequences were anaysed with the mistomaster, a web-based bioinformatics tool.

**Results:** MtDNA haplogroup U was significantly associated with susceptibility to benign and malign thyroid entities on the other hand J haplogroup was associated with a protective role for benign thyroid nodules. Besides, 8 SNPs (T146C, G185A, C194T, C295T, G16129A, T16304C, A16343G and T16362C) in mtDNA CR region were associated with the occurrence of benign and malign thyroid nodules in Turkish population. By contrast with the healthy Turkish population and HTNs, frequency of C7 repeats in D310 polycytosine sequence was found higher in cold thyroid nodules and PTC samples. Beside this, the frequency of somatic mutations in mtMSI regions including T16189C and D514 CA dinucleotide repeats were found higher in PTC samples than the benign thyroid nodules. Conversely, the frequency of somatic mutations in D310 was detected higher in HTNs than CTNs and PTCs.

**Conclusion:** mtDNA D310 instability do not play a role in tumorogenesis of the PTC but the results indicates that it might be used as a diagnostic clonal expansion biomarker for premalignant thyroid tumor cells. Beside this, D514 CA instability might be used as prognostic biomarker in PTCs. Also, we showed that somatic mutation rate is less frequent in more aggressive tumors when we examined micro- and macro carcinomas as well as *BRAF*V600E mutation.

## Introduction

Mitochondria play a central role in variety of cellular processes including metabolism, ATP production, calcium homeostasis and apoptosis (Tipirisetti et al., 2014). Shifting in glucose metabolism from OXPHOS to glycolysis has been observed in cancer cells nearly a century ago by Otto Warburg and he proposed that tumor cells, unlike the normal one, display highly glycolytic activity even in the presence of abundant oxygen for mitochondria to produce ATP by means of oxidative phosphorylation (Warburg et al., 1927). This metabolic reprogramming is known as Warburg effect (Lee and Wei, 2009). Although advantages conferred to cancer cells of the Warburg effect have not been clearly understood, it is suggested that they may relate to the ability to proliferate in the lack of oxygen, to decreased susceptibility to mitochondrial apoptosis, or to the need of the tumor cell to accumulate nutrients for increased proliferation (Lee and Wei, 2009; Nicholls and Ferguson, 2013; Su et al., 2016).

Human mitochondrial DNA (mtDNA) is a 16,569 bp double-stranded circular, maternally inherited DNA molecule. It contains 13 protein-coding genes for subunits of of the respiratory complexes CI, CIII, CIV and F-ATPase, and also encodes 22 tRNAs and 2 rRNAs. Recent studies also reported additional gene content which encodes regulatory RNAs and even additional proteins (Cobb et al., 2016; Duarte et al., 2014). Regulation and maintenance of mtDNA achieved by nuclear-encoded mitochondrial targeted proteins. Approximately 10^3^-10^4^ copies of mtDNA are present per cell. MtDNA has ~ 5-15 times higher mutation rate than the nuclear genome because of having relatively weak DNA repair mechanisms and due to its proximity to reactive oxygen species (ROS) (Payne et al., 2013). The control region (CR) of the mtDNA also including displacement loop (D-loop) extends between nucleotide positions (np) 16024-576. The D-loop which extends from around O_H_ (at the 5’ end of 7S DNA) to the termination-associated sequence (TAS) (approximately 650 nt long) has a unique triple-strand characteristic and controls the replication and transcription in the mitochondria (Nicholls and Minczuk, 2014; Sharma et al., 2005). For this reason, in literature it is assumed that mutation in the D-loop can modulate mtDNA copy number, subsequent OXPHOS dysfunction and increased ROS (De Paepe, 2012). The CR region of the mtDNA is the hotspot for both germline and somatic mtDNA alterations. In addition, the most frequent DNA alterations in the mitochondrial genome were reported in the hypervariable segment-1 (HSV1, np 16204-16383) and −2 (np 57-373) of the CR, which were mostly located in the D-loop, at various types of tumors such as head and neck, colorectal, lung, bladder, melanoma, uterine cervix and breast cancer (Ashtiani et al., 2012; Cai et al., 2011; Zhang et al., 2015). Further associations were also established between single nucleotide polymorphisms (SNPs) of the CR and various complex diseases such as; metabolic syndrome, type II diabetes mellitus (DM), neurodegenerative diseases and aging in literature (Ghezzi et al., 2005; Guney et al., 2014; Mueller et al., 2011; Tipirisetti et al., 2014; Zhang et al., 2015). But, some of the findings are still debatable. Moreover, several studies reported that mtDNA polymorphisms and mitochondrial haplogroups have either predisposing or protective role in various cancer types (Cocos et al., 2017).

Thyroid nodules are common entities, frequently diagnosed in thyroid clinics and prevalence of thyroid nodules in adult populations were detected in various studies up to 76% by using sensitive high-resolution ultrasound methods. Also, thyroid nodules are 4 times more common in women than the men because of hormonal influence like estrogen and progesterone. Iodine deficiency, age, multiparity and exposure to ionizing radiation also cause increase in nodular formation (Krohn et al., 2007; Popoveniuc and Jonklaas, 2012). Functionally, thyroid nodules have been classified as cold, warm or hot by using scintigraphy scan with Iodine^123^ or technetium ^99m^Tc pertechnetate depend on the amount of activity in the thyroid nodule compared to rest of the thyroid tissue (Gozu et al., 2004). The benign nodules, having high radioactive iodine uptake in scintigraphy represent as hot thyroid nodule (or termed as hyperfunctioning nodule) and 5% of all thyroid nodules are hot (Krohn et al., 2005). Moreover, they are very rarely malignant (approximately 1% or less) and should not be considered for fine needle aspiration (Gozu et al., 2004; Popoveniuc and Jonklaas, 2012). In spite of that the cold nodules have diminished iodine uptake and approximately 85% of all thyroid nodules are cold (or termed as hypofunctioning or isofunctioning) (Krohn et al., 2005). Furthermore, the risk of malignancy increases approximately 5% to 15% in cold thyroid nodules. Therefore, they should be considered for further clinical evaluations (Gharib and Papini, 2007).

However, thyroid cancer is the most common endocrine malignancy covers 90% of the all endocrine malignancies and it has rapidly increase in global incidence in recent decades (Cocos et al., 2017; Xing, 2013). Beside this, thyroid carcinoma was the third most common malignancy in both gender (12%) in Turkey and the second most common malignancy among Turkish women (12%) according to the 2017 Turkish health ministry official cancer report (http://kanser.gov.tr). In Turkey, the incidence of thyroid carcinoma has increased from 10.8 per 100,000 individuals in 2006 to 20.7 per 100,000 individual in 2014, representing approximately a 2-fold increase (Daglar-Aday et al., 2013).

The papillary thyroid carcinoma (PTC), which is characterized as a differentiated thyroid cancer, is the most common subtype of all thyroid cancer histological variants. PTC forms approximately 80% of the all thyroid cancers. Although, nuclear genetic alterations, underlying PTC molecular pathogenesis widely investigated and well established but the contribution of the mitochondrial genome, especially, mtDNA CR region remains unclear due to persistence of discrepancy in literature. For instance, Maximo et al. (2005) reported 40 % of the 30 adenoma, 26.3% of the 16 PTC samples had somatic mutation in D310 and 10 % of the 30 adenoma, 36.8% of the 16 PTC samples had somatic mutation in D514 whereas Ding et al. (2010) reported none of the 5 adenoma and 10.4 % of the 77 PTCsamples had somatic mutation in D310 and none of the adenoma and PTC samples had any somatic alteration in D514. In addition to that, Lohrer et al. (2002) showed only 4.2% of the 146 malign and 5 benign thyroid tumors had somatic mutation in D310. However, a few studies are interested in genomic changes of the mtDNA D-Loop region at thyroid tumors, especially PTC (Lohrer et al., 2002). Majority of the studies have been focused on mtDNA microsatellite instabilities (mtMSIs) either D310 alone or D310, D514 and np 16183-16192 altogether (Ding et al., 2010; Lohrer et al., 2002; Maximo et al., 2005; Su et al., 2016). Also, these studies have some sample size limitations. Moreover, mtDNA CR or D-loop alone was partially sequenced in these studies or the studies have not included the healthy control subjects. So, they did not considered any influence of the other SNPs rather than mtMSIs at CR of the mtDNA. But only one study in literature includes all mitochondrial genome investigation at PTC samples but the study did not have any benign thyroid tumor samples to demonstrate the influence of the mitochondrial genome in thyroid tumorigenesis (Su et al., 2016). In addition to that, Cocos et al. (2017) reported that mtDNA haplogroup K has a protective role for thyroid cancer and suggested that investigating mtDNA haplogroups is a candidate susceptibility marker for the patients with thyroid nodules.

Hence, the purpose of this study is to investigate the role of mtDNA CR variations in thyroid tumor progression and the influence of mtDNA haplogroups on susceptibility to thyroid tumors.

## Materials & Methods

### Patients

Totally 52 toxic multinodular guatr (MNG), 53 MNG, 48 PTC patients and 104 healthy control subjects without any evidence of thyroid disorder were enrolled to the study. The study approved by the local ethics committee. A total of consecutive 108 hot thyroid nodules (HTNs), 95 cold thyroid nodules (CTNs) and 105 surrounding tissues obtained from unrelated patients who underwent subtotal or near-total thyroidectomy for toxic and nontoxic MNG. All 108 HTNs and 95 CTNs were identified by ultrasound and scinthigraphy between 2000-2004 at Marmara University, School of Medicine and Kartal Dr. Lütfi Kırdar Training and Education Hospital Pathology as described previously (Gozu et al., 2005; Gozu et al., 2006). All preoperatively identified nodules were also characterized during surgery and postoperatively by histology according to the WHO criteria (Eszlinger et al., 2006). The 48 PTC samples and surrounding tissues were obtained from pathology archive materials, and were diagnosed histopathologically according to the WHO criteria revealed in 2004 (Vuong et al., 2016). PTC samples were divided into two groups according to the tumor diameter. Samples with tumor diameter ≤ 1cm were classified as micro-PTCs and the others with tumor diameter >1cm were classified as macro-PTCs. 21 samples were classified as micro-PTCand 27 samples were classified as macro-PTC. Cell subtype determinations of the 48 PTC patients were performed in formalin fixed paraffin embedded tissue (FFPE) samples. For this purpose the FFPE samples were stained with hematoxylene-eosine and analyzed by light microscopy. Clinical cell subtypes of the PTC samples were indicated in Supplementary Table 4.

**Table 4.** Summary of the somatic mtDNA control region MSI alterations in 203 benign nodules and 48 papillary thyroid tumors.

| Nodule no | Nodule type | mtDNA CR status and somatic alterations |  |  |  |
| --- | --- | --- | --- | --- | --- |
|  |  | T16189C | D310* | D514** | D568***<br>(6C→9C/10C) |
| 1.1 | HTN<sup>a</sup> | T | C8/C9→C8/C9/C10 | 5CA | 6C |
| 1.2 | HTN | T | C8/C9→C8/C9/C10 | 5CA | 6C |
| 2 | HTN | C | C8→C8/C9 | 5CA | 6C |
| 3 | HTN | T | C8→C8/C9 | 5CA | 6C |
| 4 | HTN | T | C8→C7/C8 | 5CA | 6C |
| 5 | HTN | T | C7→C8/C9 | 4CA | 6C→10C |
| 6 | HTN | T | C8→C8/C9 | 6CA | 6C |
| 7 | HTN | T→C | C8→C8/C9 | 5CA | 6C |
| 8 | HTN | T | C7→C7/C8/C9 | 5CA | 6C |
| 9.1 | HTN | T | C9/C10→C8/C9 | 5CA | 6C |
| 9.2 | HTN | T | C9/C10→C8/C9 | 5CA | 6C |
| 9.3 | HTN | T | C9/C10→C8/C9 | 5CA | 6C |
| 10.1 | HTN | C | C7→C7/C8 | 5CA | 6C |
| 10.2 | HTN | C | C7→C7/C8 | 5CA | 6C |
| 11.1 | HTN | T | C9→C8 | 7CA | 6C |
| 11.2 | HTN | T | C9→C8 | 7CA | 6C |
| 12.1 | HTN | T | C7 | 5CA | 6C→10C |
| 12.2 | HTN | T | C7→C8 | 5CA | 6C→10C |
| 13 | HTN | T | C7 | 5CA | 6C→10C |
| 14 | HTN | T | C7→C7/C8 | 5CA | 6C |
| 15 | HTN | T | C8→C8/C9 | 5CA | 6C |
| 16 | HTN | T | C8/C9→C9/C10 | 5CA | 10C |
| 17 | HTN | T | C8→C9/C10 | 5CA | 6C |
| 18.1 | HTN | C | C8→C8/C9 | 5CA | 6C |
| 18.2 | HTN | C | C8→C8/C9 | 5CA | 6C |
| 19 | HTN | T | C8→C8/C9 | 5CA | 10C |
| 1 | CTN<sup>b</sup> | C | C7 | 5CA→5CA/6CA | 6C |
| 2.1 | CTN | T→C | C7→C7/C8/C9 | 5CA | 6C |
| 2.2 | CTN | T→C | C7→C8/C9 | 5CA | 6C |
| 3 | CTN | T→C | C8→C7/C8 | 5CA | 6C |
| 4.1 | CTN | T | C8→C8/C9 | 5CA | 6C |
| 4.2 | CTN | T | C8→C8/C9 | 5CA | 6C |
| 5 | CTN | T | C8→C8/C9 | 5CA | 6C |
| 6 | CTN | T | C7 | 7CA→6CA/7CA | 6C |
| 7.1 | CTN | T | C8 | 5CA→4CA/5CA | 6C |
| 7.2 | CTN | T | C8 | 5CA→4CA/5CA | 6C |
| 8 | CTN | T→C | C7 | 5CA | 6C |
| 9.1 | CTN | T | C7/C8→C7 | 5CA | 6C |
| 9.2 | CTN | T | C7/C8→C7 | 5CA | 6C |
| 9.3 | CTN | T | C7/C8→C7 | 5CA | 6C |
| 9.4 | CTN | T | C7/C8→C7 | 5CA | 6C |
| 10 | CTN | C | C8 | 7CA→6CA/7CA | 6C |
| 1 | PTCCV<sup>c</sup> | T | C8 | 5CA→4CA | 6C |
| 2 | PTCCV | T→C | C7/C8 | 4CA/5CA | 6C |
| 3 | PTCCV | T→C | C7 | 4CA/5CA | 6C |
| 4 | PTCCV | T | C8/C9→C7/C8/C9 | 5CA→4CA | 6C |
| 5 | PTMCCV<sup>d</sup> | T | C7 | 5CA→4CA | 10C |
| 6 | PTMCCV | T→C | C8/C9 | 5CA | 6C |
| 7 | PTMCCV | T→C | C8/C9 | 5CA | 10C |
| 8 | PTMCCV | T | C7/C8 | 5CA→4CA | 6C |
| 9 | PTMCCV | T→C | C7 | 5CA→4CA | 6C |
| 10 | PTMCCV | T | C8/C9→C7/C8 | 5CA→4CA | 6C |
| 11 | PTMCCV | T | C8→C7/C8 | 5CA→4CA | 6C |
| 12 | PTCFV<sup>e</sup> | T | C8→C7/C8 | 5CA→4CA | 6C |
| 13 | PTMCFV<sup>f</sup> | T | C7/C8 | 5CA→4CA | 6C |
| 14 | PTMCFV | T→C | C7/C8 | 4CA/5CA | 6C |
| 15 | PTCOV<sup>g</sup> | T | C7 | 5CA→4CA | 6C |
| 16 | PTCOV | T | C7/C8 | 5CA→4CA | 6C |
| 17 | PTMCOV<sup>h</sup> | T→C | C7→C7/C8 | 5CA→4CA | 6C |
|  |  | HTNs<br>n(%) | CTNs<br>n(%) | PTCs<sup>i</sup><br>n(%) | *p* value<sup>j</sup> |
| Nodular tissue harboring<br>at least one somatic<br>mutation in mtMSIs |  | 26(24.1) | 16(16.8) | 17(35.4) | **<u>0.046</u>** |
| Nodular tissue harboring<br>at least two somatic<br>mutation in mtMSIs |  | 5(4.6) | 3(3.2) | 6(12.6) | 0.061 |
| Nodular tissue harboring<br>at least two somatic<br>mutation in mtMSIs |  | **5(4.6)** | **3(3.2)** | **4(19.0)<sup>k</sup>** | **<u>0.012</u>** |

\* 8C in poly-C stretch at D310 is the common genotype in Turkish population.
\*\* 5CA dinucleotide repeat is the wild-type genotype.
\*\*\* 6C in poly-C stretch at D568 is the wild-type genotype.
<sup>a</sup>HTN :Hot thyroid nodule
<sup>b</sup>CTN :Cold thyroid nodule
<sup>c</sup> PTCCV :Papillary thyroid carcinoma classical variant
<sup>d</sup> PTMCCV :Papillary thyroid microcarcinoma classical variant
<sup>e</sup> PTCFV :Papillary thyroid carcinoma follicular variant
<sup>f</sup> PTMCFV :Papillary thyroid microcarcinoma follicular variant
<sup>g</sup> PTCOV :Papillary thyroid carcinoma oncocytic variant
<sup>h</sup> PTMCOV :Papillary thyroid microcarcinoma oncocytic variant
<sup>i</sup>PTCs :Papillary thyroid carcinomas
<sup>j</sup> :Pearson's chi-square test
<sup>k</sup> :Data belongs to the microPTC group

All healthy control subjects had no evidence of thyroid autoimmune or non-autoimmune disease. They are also subjected a careful investigation of personal and family history, clinical examination, thyroid function testing, thyroid autoantibody testing, and thyroid ultrasound testing.

### DNA Isolation

Genomic DNA was isolated from frozen CTN and HTN tissue specimens using standard techniques as described previously (Gozu et al., 2005; Gozu et al., 2006). Genomic DNA from FFPE archive material of the PTC tissue samples was isolated by using QIAampDNA FFPE tissue kit (Qiagen, USA) according to the manufacturer’s instructions. Following the DNA isolation, until the start of the study DNAs were archived at −20°C in deep freezer.

### Polymerase Chain Reaction (PCR) and DNA Sequencing

mtDNA CR region (np16024-576) and tRNA^phe^ gene (np577-647) were amplified entirely and 12S rRNA gene (np647-921) was amplified partially using the primers and conditions as described previously (Levin et al., 1999). List of the primers used were given in the Supplementary Table 1. However, in FFPE samples from PTC patients, the target sequence was amplified by using Hotstar Master Mix (Qiagen, USA) according to the manufacturer’s manual and mtDNA D-loop region between 16301-134 nt. was amplified by hot start PCR with using newly designed primers as mentioned in Supplementary Table 1. The PCR products were sequenced utilizing DTCS quick start kit (Beckman Coulter, USA). The sequencing reaction was carried out in a Proflex thermocycler at 96°C for 20s, 50°C for 20s and 60°C for 4 minutes according to the manufacturer’s manual. Sequence analysis was performed on the automatic DNA sequencer (Beckman Coulter GenomeLabGeXP Genetic Analysis System, USA). DNA sequences and chromatograms obtained were examined by using GenomeLabGeXP Genetic Analysis System Version 10.2 DNA sequencing program (Beckman Coulter, USA).

**Table 1.** Influence of mtSNPs on benign and malign thyroid lesions.

| T146C polymorphism |  |  |  |  | T152C polymorphism |  |  |  |  |
| --- | --- | --- | --- | --- | --- | --- | --- | --- | --- |
| Group | 146T | 146C | *p* value<sup>a</sup> | OR (95% CI) | Group | 152T | 152C | *p* value | OR (95% CI) |
| HTN(n=65) | 59(90.8) | 6(9.2) | 0.189 | 1.921 (0.716 - 5.159) | HTN(n=65) | 49(75.4) | 16(24.6) | 0.845 | 1.074 (0.526 - 2.194) |
| CTN(n=56) | 53(94.6) | 3(5.4) | **0.045** | 3.452 (0.966 - 12.342) | CTN(n=56) | **48(85.7)** | **8(14.3)<sup>b</sup>** | 0.088 | 2.104 (0.884 - 5.009) |
| PTC(n=48) | 47(97.9) | 1(2.1) | **0.011** | 9.184 (1.185 - 71.180) | PTC(n=48) | **29(60.4)** | **19(39.6)<sup>b</sup>** | 0.089 | 0.535 (0.259 - 1.106) |
| NC(n=104) | 87(83.7) | 17(16.3) |  |  | NC(n=104) | 77(74.0) | 27(26.0) |  |  |
| G185A polymorphism |  |  |  |  | C194T polymorphism |  |  |  |  |
| Group | 185G | 185A | *p* value | OR (95% CI) | Group | 194C | 194T | *p* value | OR (95% CI) |
| HTN(n=65) | 64(98.5) | 1(1.5) | **0.026** | 7.570 (0.954 - 60.093) | HTN(n=65) | 63(95.4) | 3(4.6) | 0.807 | 0.840 (0.182 - 3.878) |
| CTN(n=56) | 54(96.4) | 2(3.6) | 0.122 | 3.194 (0.682 - 14.949) | CTN(n=56) | 54(96.4) | 2(3.6) | 0.930 | 1.080 (0.192 - 6.088) |
| PTC(n=48) | 47(97.9) | 1(2.1) | 0.071 | 5.559 (0.697 - 44.362) | PTC(n=48) | 42(87.5) | 6(12.5) | **0.045** | 0.280 (0.075 - 1.043) |
| NC(n=104) | 93(89.4) | 11(10.6) |  |  | NC(n=104) | 100(96.2) | 4(3.8) |  |  |
| T195C polymorphism |  |  |  |  | C295T polymorphisms |  |  |  |  |
| Group | 195T | 195C | *p* value | OR (95% CI) | Group | 295C | 295T | *p* value | OR (95% CI) |
| HTN(n=65) | 53(81.5) | 12(18.5) | 0.782 | 1.117 (0.508 - 2.459) | HTN(n=65) | 63(96.9) | 2(3.1) | **0.036** | 4.500 (0.981 - 20.636) |
| CTN(n=56) | 47(83.9) | 9(16.1) | 0.524 | 1.321 (0.560 - 3.119) | CTN(n=56) | 55(98.2) | 1(1.8) | **0.022** | 7.857 (1.000 - 61.728) |
| PTC(n=48) | 33(68.8) | 15(32.2) | 0.136 | 0.557 (0.256 - 1.209) | PTC(n=48) | 43(89.6) | 5(10.4) | 0.712 | 1.229 (0.412 - 3.666) |
| NC(n=104) | 83(79.8) | 21(20.2) |  |  | NC(n=104) | 91(87.5) | 13(12.5) |  |  |
| D568 (insCCCC or insCCCCC) polymorphism |  |  |  |  | G709A polymorphism |  |  |  |  |
| Group | 573C | insCCCC or insCCCCC | *p* value | OR (95% CI) | Group | 709G | 709A | *p* value | OR (95% CI) |
| HTN(n=65) | 61(93.8) | 4(6.2) | 0.299 | 0.453 (0.098 - 2.093) | HTN(n=65) | 47(72.3) | 18(27.7) | 0.109 | 0.547 (0.260 - 1.150) |
| CTN(n=56) | 54(96.4) | 2(3.6) | 0.812 | 0.802 (0.130 - 4.947) | CTN(n=56) | 49(87.5) | **7(12.5)<sup>b</sup>** | 0.424 | 1.465 (0.572 - 3.754) |
| PTC(n=48) | 45(93.7) | 3(6.3) | 0.322 | 0.446 (0.087 - 2.293) | PTC(n=48) | 34(70.8) | **14(29.2)<sup>b</sup>** | 0.096 | 0.508 (0.228 - 1.135) |
| NC(n=104) | 101(97.1) | 3(2.9) |  |  | NC(n=104) | 86(82.3) | 18(17.3) |  |  |
| G16129A polymorphism |  |  |  |  | T16189C polymorphism |  |  |  |  |
| Group | 16129G | 16129A | *p* value | OR (95% CI) | Group | 16189T | 16189C | *p* value | OR (95% CI) |
| HTN(n=65) | 58(89.2) | 7(10.8) | 0.235 | 0.507 (0.163 - 1.583) | HTN(n=65) | 56(86.2) | 9(13.8) | 0.550 | 1.302 (0.547 - 3.102) |
| CTN(n=56) | 51(91.1) | 5(8.9) | 0.451 | 0.624 (0.182 - 2.145) | CTN(n=56) | 47(83.9) | 9(16.1) | 0.842 | 1.093 (0.455 - 2.624) |
| PTC(n=48) | 35(68.8) | 13(31.2) | **0.0002** | 0.165 (0.058 - 0.467) | PTC(n=48) | 38(79.2) | 10(20.8) | 0.602 | 0.795 (0.336 - 1.884) |
| NC(n=104) | 98(87.9) | 6(5.8) |  |  | NC(n=104) | 86(82.7) | 18(17.3) |  |  |
| C16292T polymorphism |  |  |  |  | C16296T polymorphism |  |  |  |  |
| Group | 16292C | 16292T | *p* value | OR (95% CI) | Group | 16296C | 16296T | *p* value | OR (95% CI) |
| HTN(n=65) | 60(92.3) | 5(7.7) | 0.067 | 0.235 (0.044 - 1.251) | HTN(n=65) | 59(90.8) | **6(9.2)<sup>c</sup>** | 0.149 | 0.393 (0.107 - 1.451) |
| CTN(n=56) | 51(91.1) | 5(8.9) | **0.039** | 0.200 (0.038 - 1.067) | CTN(n=56) | 54(96.4) | 2(3.6) | 0.930 | 1.080 (0.192 - 6.088) |
| PTC(n=48) | 48(100) | 0(0) | 0.333 | 1.020 (0.993 - 1.047) | PTC(n=48) | 48(100) | **0(0)<sup>c</sup>** | 0.169 | 1.040 (1.001 - 1.081) |
| NC(n=104) | 102(98.1) | 2(1.9) |  |  | NC(n=104) | 100(96.2) | 4(3.8) |  |  |
| T16304C polymorphism |  |  |  |  | A16343G polymorphism |  |  |  |  |
| Group | 16304T | 16304C | *p* value | OR (95% CI) | Group | 16343A | 16343G | *p* value | OR (95% CI) |
| HTN(n=65) | 56(86.2) | 9(13.8) | **0.018** | 0.249 (0.073 - 0.845) | HTN(n=65) | 59(90.8) | 6(9.2) | **0.030** | 0.193 (0.038 - 0.986) |
| CTN(n=56) | 52(92.9) | 4(7.1) | 0.361 | 0.520 (0.125 - 2.164) | CTN(n=56) | 52(92.9) | 4(7.1) | 0.097 | 0.255 (0.045 - 1.438) |
| PTC(n=48) | 46(95.8) | 2(4.2) | 0.925 | 0.920 (0.163 - 5.205) | PTC(n=48) | 45(93.7) | 3(6.3) | 0.164 | 0.294 (0.047 - 1.821) |
| NC(n=104) | 100(96.2) | 4(3.8) |  |  | NC(n=104) | 102(98.1) | 2(1.9) |  |  |
| C16355T polymorphism |  |  |  |  | T16362C polymorphism |  |  |  |  |
| Group | 16355C | 16355T | *p* value | OR (95% CI) | Group | 16362T | 16362C | *p* value | OR (95% CI) |
| HTN(n=65) | 60(92.3) | 5(7.7) | 0.439 | 0.606 (0.168 - 2.181) | HTN(n=65) | 64(98.5) | 1(1.5) | 0.389 | 2.560 (0.280 - 23.482) |
| CTN(n=56) | 51(91.1) | **5(8.9)<sup>b</sup>** | 0.304 | 0.515 (0.143 - 1.862) | CTN(n=56) | 49(87.5) | 7(12.5) | **0.039** | 0.280 (0.078 - 1.002) |
| PTC(n=48) | 48(100) | **0(0)<sup>b</sup>** | 0.122 | 1.051 (1.006 - 1.097) | PTC(n=48) | 46(95.8) | 2(4.2) | 0.925 | 0.920 (0.163 - 5.205) |
| NC(n=104) | 99(95.2) | 5(4.8) |  |  | NC(n=104) | 100(96.2) | 4(3.8) |  |  |
| A16399G polymorphism |  |  |  |  |  |  |  |  |  |
| Group | 16399A | 16399G | *p* value | OR (95% CI) |  |  |  |  |  |
| HTN(n=65) | 62(96.9) | 3(3.1) | 0.554 | 0.614 (0.120 - 3.137) |  |  |  |  |  |
| CTN(n=56) | 51(91.1) | **5(8.9)<sup>b</sup>** | 0.094 | 0.303 (0.070 - 1.318) |  |  |  |  |  |
| PTC(n=48) | 48(100) | **0(0)<sup>b</sup>** | 0.235 | 1.030 (0.996 - 1.064) |  |  |  |  |  |
| NC(n=104) | 101(97.1) | 3(2.9) |  |  |  |  |  |  |  |

CTN: Cold thyroid nodules
HTN: Hot thyroid nodules
PTC: Papillary thyroid carcinomas
NC: Healthy Control
<sup>a</sup> Pearson's chi square correlation tests
<sup>b</sup> Indicates statistically significant difference between CTN and PTC patient groups, which data was not shown.
<sup>c</sup> Indicates statistically significant difference between HTN and PTC patient groups, which data was not shown.

### Sequence Analysis and Haplogroup Classification

The entire sequences of mtDNA CR and tRNA^phe^ and 12S rRNA coding gene region were aligned to the latest revised Cambridge Reference Sequence (rCRS) (Genebank accession number: NC_012920.1) by BLAST (http://www.ncbi.nlm.nih.gov/blast). In order to identify the mtDNA polymorphisms, all of the sequences were analyzed for the variation in sequence by using Mito Tool Programme. Also, the sequences subsequently were analyzed in Mitomaster, a web-based bioinformatics program, to verify the results and confirm the polymorphisms (http://www.mitomap.org/foswiki/bin/view/MITOMASTER/WebHome). The polymorphisms not recorded in the Mitomap database were regarded as novel. Also, the somatic mutations were defined as difference in mtDNA sequence between nodular/tumor tissues and surrounding tissues. The haplogroup prediction from partial mtDNA sequence was made by using Mitomaster program. This program uses HaploGrep 2 with Phylotree 17 for haplogroup determination.

### Statistical Analysis

The SPSS 15.0 software (SPSS Inc., Chicago, IL, USA) package was used for the statistical analyses. To compare frequency distribution of the polymorphisms at mtMSIs and non-mtMSIs in each group and prevalence of somatic mutations in each group, the Pearson’s chi-square test was used. The Pearson’s chi-square test was used also to evaluate the haplogroup distribution among the groups. The results were evaluated with 95% CIs and *p* value <0.05 was considered as significant.

## Results

### mtDNA Polymorphisms and Haplogroups

A total of 108 HTNs, 95 CTNs and their surrounding tissues belong to 52 toxic MNG and 53 non-toxic MNG patients included in this study. Also samples from 48 PTC patients were enrolled the study. 267 different sequence variants were detected in 251 nodular,153 surrounding tissues and 104 healthy control samples compared with rCRS (Supplementary Table 2). Overall, 163, 154, 136 and 168 different sequence variants were observed in HTN, CTN, PTC and healthy control samples, respectively. Seventeen novel polymorphisms and one novel somatic mutation were detected in this study. Eight of them were transversions and 10 were them transitions (Supplementary Table 2)

**Table 2.** Comparison of common haplogroup distribution in the study groups versus healthy controls.

| H haplogroup |  |  |
| --- | --- | --- |
| Group | Haplogroup frequency n(%) | *p* value |
| HTN (n=65) | 19(29.23) | 0.894 |
| CTN (n=56) | 18(32.14) | 0.802 |
| PTC (n=48) | 14(29.16) | 0.863 |
| NC (n=104) | 31(29.81) |  |
| HV haplogroup |  |  |
| Group | Haplogroup frequency n(%) | *p* value |
| HTN (n=65) | 3(4.61) | 0.246 |
| CTN (n=56) | 3(5.35) | 0.362 |
| PTC (n=48) | 1(2.08) | **0.056** |
| NC (n=104) | 10(9.62) |  |
| J haplogroup |  |  |
| Group | Haplogroup frequency n(%) | *p* value |
| HTN (n=65) | 2(3.07) | **0.039** |
| CTN (n=56) | 1(1.8) | **0.022** |
| PTC (n=48) | 5(10.42) | 0.598 |
| NC (n=104) | 13(12.50) |  |
| T haplogroup |  |  |
| Group | Haplogroup frequency n(%) | *p* value |
| HTN (n=65) | 11(16.92) | 0.097 |
| CTN (n=56) | 5(8.92) | 0.926 |
| PTC (n=48) | 4(8.32) | 0.948 |
| NC (n=104) | 9(8.65) |  |
| U haplogroup |  |  |
| Group | Haplogroup frequency n(%) | *p* value |
| HTN (n=65) | 14(21.54) | **0.027** |
| CTN (n=56) | 12(21.42) | **0.007** |
| PTC (n=48) | 10(20.80) | **0.058** |
| NC (n=104) | 10(9.62) |  |

Nevertheless, 29 sequence variations have high frequency among 267 different sequence variations. The frequency of polymorphisms when the HTN, CTN, and PTC groups compared with healthy control group, 8 of these SNPs (T146C, G185A, C194T, C295T, G16129A, T16304C, A16343G and T16362C) are statistically significant (*p*<0.05) (Table 1). The T→C transition at np 194 and the G→A transition at np16129 increased in PTCs compared to healthy control subjects those indicates susceptibility to malign transformation (*p*=0.045 and 0.0002, respectively). On contrary, the frequency of T146C decreased in benign and malign thyroid tumors that reveals a protective role (Table 1). Additionally, although T195C frequency is higher in PTC group (32.2%) than the other groups, it is not statistically significant (*p*=0.136). High frequency of T→C transition at np 16304 and A→G transition at np 16343 indicates increased risk of HTN formation (*p*=0.018 and 0.030, respectively). Beside this, C→T transition at np 295 significantly lower in benign thyroid nodules than the healthy control subjects (*p*=0.036 and 0.022, respectively). Increased frequency of T→C transition at np16362 signs decreased risk of malignancy at CTN nodules compared to the PTCs. However, polymorphisms at nps 152, 709, 16296, 16355 and 16399 were not statistically different between patients and healthy control subjects but the frequencies of T152C and G709A significantly higher in PTCs than CTNs which indicates a susceptibility to malign transformation in CTNs. On the other hand, the frequencies C16355T and A16399G significantly higher in CTNs than PTCs which points out a protective role. Meanwhile, high frequency of C16296T in HTNs than PTCs may indicate a protective role also.

The haplogroup of the patients were predicted by using Mitomaster from the partial sequence of the mtDNA (np16011-921) and summarized in Supplementary Table 3. H, HV, J, T and U are the most common haplogroups found in this study. Frequency of these 5 haplogroups was compared with the healthy control subjects by using Pearson’s chi-square test (Table 2). Frequencies of H and T haplogroup distribution are similar among benign and malign thyroid patients and healthy control groups. Frequency of the J haplogroup in patients carrying HTNs and CTNs are lower than healthy control group (*p*=0.039 and 0.022, respectively) (Table 2). Frequency of the HV haplogroup in PTC patients is lower than healthy control group but not statistically significant (*p*=0.056). Frequency of the U haplogroup is significantly higher in patients with benign thyroid nodules (HTNs and CTNs groups) compared with the healthy control subjects (*p*=0.027 and 0.007, respectively). However, frequency of U haplogroup is also higher in PTC group than the healthy control subjects but not statistically significant (*p*=0.058) (Table 2).

**Table 3.** Frequency of somatic mutations in mtMSIs and non-mtMSI sequence in mtDNA control region.

|  | HTN<br>Total 108<br>n(%) | CTN<br>Total 95<br>n(%) | PTC<br>Total 48<br>n(%) | *p* value |
| --- | --- | --- | --- | --- |
| Somatic D310 mutations | **<u>24 (22.2)</u>** | **<u>10 (10.4)</u>** | **<u>5 (10.4)</u>** | **<u>0.04</u>** |
| Somatic T16189C mutations | **<u>1 (0.9)</u>** | **<u>3 (3.2)</u>** | **<u>7 (14.58)<sup>a</sup></u>** | **<u>0.001</u>** |
|  |  |  | 2 (7.4)<sup>b</sup> | 0.152 |
|  |  |  | **<u>5 (23.8)<sup>c</sup></u>** | **<u><0.00001</u>** |
| Somatic T16189C mutations without any accompanied polymorphism in np16184-16193 | **<u>1(0.9)</u>** | **<u>1 (1.1)</u>** | **<u>5 (10.4)<sup>a</sup></u>** | **<u>0.002<sup>d</sup></u>** |
| Somatic D514CA mutations | **<u>1 (0.9)</u>** | **<u>4 (4.2)</u>** | **<u>12 (25.00)</u>** | **<u>0.0003</u>** |
| Somatic D568 mutations | 3 (2.7) | 0 (0) | 1 (2.08) | 0.276 |
| Somatic mutations in nonmtMSI sequence | **<u>5 (4.6)</u>** | **<u>4 (4.2)</u>** | **<u>8 (16.7)<sup>a</sup></u>** | **<u>0.01</u>** |
|  |  |  | 3 (11.1)<sup>b</sup> | 0.338 |
|  |  |  | **<u>5 (23.8)<sup>c</sup></u>** | **<u>0.002</u>** |

<sup>a</sup> represents all PTC samples.
<sup>b</sup> represents macro-PTC samples.
<sup>c</sup> represents micro-PTC samples.
<sup>d</sup> In this statistical calculation , 5 mutated samples in PTC group composed of 1 macro-carcinoma and 4 micro-carcinoma samples. Prevalence of T16189C somatic mutation alone at nodules in this group is 19% (4/21). If microcarcinoma samples compared with the other groups, obtained result is more significant ( $p= 6.7 \times 10^{-6}$ )

### MtDNA microsatellite instabilities (mtMSIs) in the control region D310

The D310 is a highly homopolymeric C stretch in the D-loop region between np 303 and 315. It is a highly electrophilic and mutational hot spot region in primary tumors. Also it is considered as genetic marker to identify the tumor progression in various tumors (Ding et al., 2010). Twenty-four of the 108 HTNs (22.2%), 10/95 (10.4%) of the CTNs and 5/48 (10.4%) of the PTC samples were harbored with somatic mutations in D310 region in this study (Table 3). The mutations were C insertions varied from 1 to 4 and summarized in the Table 4, Supplementary Tables 2 and 4. Also, sample chromatograms for each variant was shown in Supplementary Figure 1. The frequency of somatic mutations is significantly higher in HTN group than the others (*p*=0.04) (Table 3). The low D310 somatic mutation frequency in CTNs and PTCs indicates that somatic mutations occurred in early stages of tumorigenesis and could not associated with the high risk of papillary thyroid cancer development.

**Figure 1.**
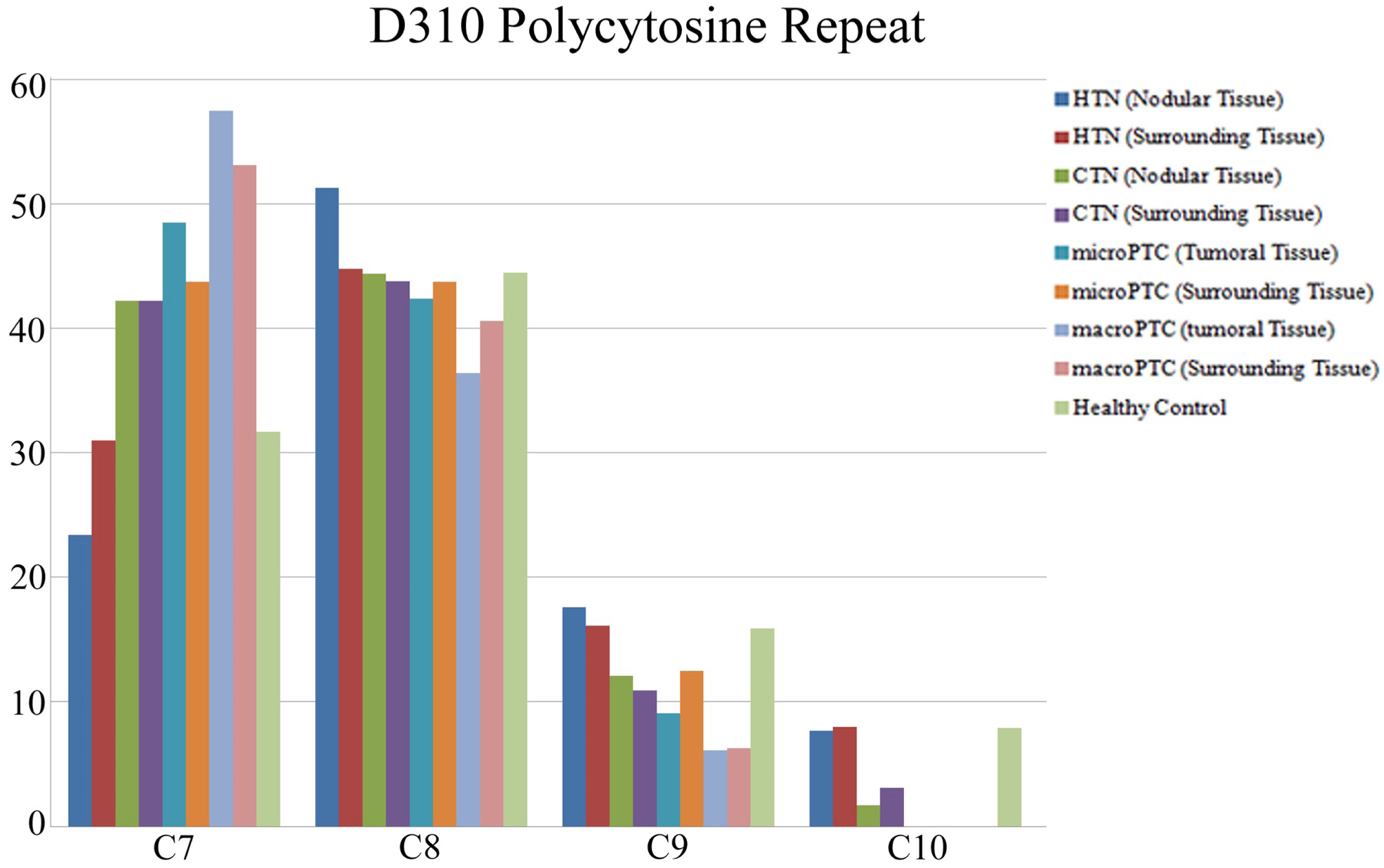
Distribution of poly-C stretch at D310 in benign thyroid nodules (CTNs (n=95) and HTNs (n=108)), malign thyroid nodules (micro-PTCs (n=21) and macro-PTCs (n=27)), their surrounding tissues (n=153) and healthy control subjects (n=104). Frequency of C7 repeat significantly higher in macro-PTC samples (*p*=0.01). Multiple sequences (heteroplasmy) may be found in one tissue sample.

Beside this, polymorphisms between np303-309 at D310 region in a normal human population consist of variations in the number of cytosine repeats, which range most commonly between seven to nine cytosine repeats (Xu et al., 2012). In this study, the poly-C stretch at D310 C7TC6 to C10TC6 refers to C7 to C10 according to the poly-C repeats. In the healthy Turkish population, it is reported that C7 frequency is in the range of 34.1 to 37.93% while C8, C9 and C10 frequencies change between 62.8 to 65.9% (Aral et al., 2006; Yacoubi-Loueslati et al., 2009). In this study, C7 repeat was found 31.7% in healthy control group, on the other hand C8, C9, C10 repeats were found to be 44.5%, 15.9% and 7.9%, respectively (Figure 1). So, the C8 repeat is the most common motif in Turkish population while the C7, C9 and C10 repeats are the rare. In this study, C8, C9 and C10 frequencies were higher in HTNs (51.3, 17.6 and 7.7%) than the CTNs (44.4, 12.1 and 1.7%) and PTCs (40.9, 6.1 and 0.0% respectively). On the other hand, C7 frequency is more prevalent in all PTC samples (53%) than the HTNs (23.4%), the CTNs (42.2) and the normal population value (31.7 %) (*p*=0.003) (data has not shown). Moreover, when the distribution of poly-C repeat in micro- and macro-PTCs compared with HTNs, CTNs and healthy control subjects, C7 frequency dramatically increase from HTNs (23.4%) to macro-PTCs group (57.5%) through healthy control subjects (31.7%), CTNs (42.2%) and micro-PTCs group (48.5) and C7 frequency was significantly higher in macro-PTCs than the other groups (*p*=0.01) (Figure 1). Moreover, C7 repeat is reached the highest frequency (66.7%) in *BRAF*V600E(+) PTC patients and in this *BRAF*V600E (+) patient group, 3 of them had both lymph node metastasis and C7 motif (Supplementary Table 5). Although C10 exists in HTNs (7.7%) similar to healthy control subjects (7.9%), but it is rare in CTNs (1.7%) and completely disappeared in all PTC samples (0.0%). Also C9 repeat frequency in HTNs (17.6%) is similar to the healthy control subjects (15.9%) but C9 frequency is proportionally decreased in CTNs, micro-PTCs, macro-PTCs group (12.1, 9.1 and 6.1%) (Figure 1) and completely disappeared in *BRAF*V600E(+) samples (Supplementary Table 5). At first glance, it is obviously seen that the frequencies of C7, C8, C9 and C10 repeats reported in healthy Turkish population diagrammatically turn upside down from CTNs, to micro- and macro-PTCs. It seems that increase in C7 repeat to 57.5% indicates selective clonal expansion in macro-PTCs on the other hand, decrease in C7 frequency to 23.4% might be an inducer for HTN formation. In other words, combination with the other molecular mechanisms in nucleus like activating TSHR mutations causing thyroid autonomy, the thyroid cells having C8 poly-C repeat or more might propagate the HTNs, in which malign thyroid transformation has been seen very rare.

### D514 CA

Five CA dinucleotide repeats are located in the CR between np514 and 524 which previously reported as a microsatellite instability region (Sharma et al., 2005). 5CA repeat is the most common genotype in normal populations. In healthy Turkish population, we found 5CA repeat frequency as 85.6% whereas 4CA, 6CA, 7CA and 8CA repeats were found as 7.7%; 2.9%; 2.9% and 0.9%, respectively (Figure 2). Somatic mutations in D514 were detected 0.9% (1/108) in HTNs, 4.2% (4/95) in CTNs and 25% (12/48) in PTCs in this study (*p*=0.0003) (Table 3). On the other hand, D514 CA repeat frequencies compared among the groups, 4CA repeats in PTCs is 2.6-fold higher than HTNs and 4.5-fold higher than CTNs (*p*<0.0001; data has not shown). 6CA, 7CA and 8CA repeats were detected fewer in our study groups. 6CA repeats were found in 4 HTNs, 8 CTNs and 2 PTCs. 7 CA repeats were assessed in 10 HTNs, 2 CTNs and 1 PTCs whereas only 1 PTC and healthy control samples were detected having 8CA repeats. If the statistical analysis were re-calculated splitting PTC group micro- and macro-carcinomas, prevalence of 4CA repeats among the groups is much more statistically significant (*p*=0.00001). The prevalence of 4CA repeats in micro-PTC is 37.3% and 35.3% in macro-PTC. However 6CA, 7CA and 8CA repeats in D514 were found in micro-PTC as 7.2%, 3.7% and 3.7 respectively, whereas no macro-PTC has one yet (Figure 2). From the obtained data, it could be concluded that either the high somatic mutation frequency or 4CA repeat frequency at D514 in both micro- or macro-papillary thyroid carcinomas might be associated with tumor progression in the thyroid. D514 CA repeat distribution in PTC group is summarized in Supplementary Table 4.

**Figure 2.**
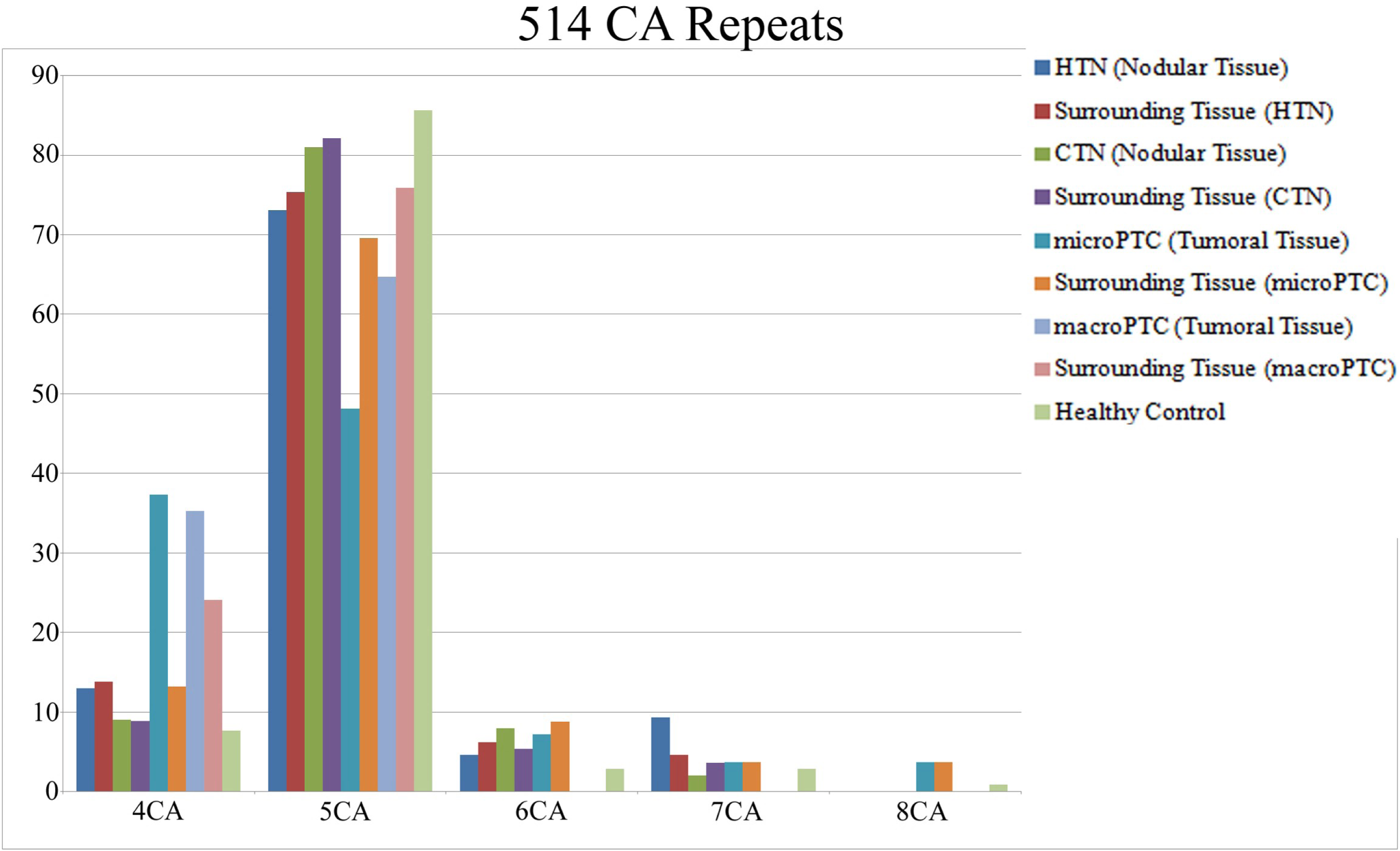
Distribution of CA repeats at D514 in benign thyroid nodules (CTNs (n=95) and HTNs (n=108)), malign thyroid nodules (micro-PTCs (n=21) and macro-PTCs (n=27)), their surrounding tissues (n=153) and healthy control subjects (n=104). Frequency of 4CA repeats significantly higher in PTC samples (*p*=0.00001). Multiple sequences (heteroplasmy) may be found in one tissue sample.

### D568

Another poly-C stretch, composed of 6 cytosines, is located between np568-573 in the mtDNA CR. Rare polymorphisms like insertion of cytosine in this poly-C stretch were reported in thyroid cancers (Maximo et al., 2005). Of the 108 nodular tissues in the HTNs, 4/108 (3.7%) and 1/48 (2.08%) of the PTC samples has somatic mutation in D568 whereas no CTNs has one (*p*=0.276) (Table 3).

### T16189C (poly-C stretch formation between np 16184-16193)

The other poly-C stretch, which is 10 base pair length or more, is occurred if there is thymine/cytosine base substitution in np16189 at mtDNA D-loop region and reported as mtMSI and mutational hotspot region previously. A number of literatures designated an association between T16189C polymorphism and disease such as, type 2 diabetes, cardiopathy, endometrial cancer risk, metabolic syndrome and melanoma. Also, T16189C polymorphism is associated with changes in mtDNA copy number in literature. Beside this, it is claimed that any accompanied polymorphism such as C16186T, C16188T or C16192T diminishes the poly-C stretch formation in this region and it might be neutralize the effect of T16189C polymorphism (Mueller et al., 2011). For this reason, we calculated and analyzed statistically both prevalence of T16189C polymorphism and poly-C stretch separately by Pearson’s chi square test and less than 10 base pair length was excluded. All the different variants between np16184 and 16193 were shown in Supplementary Figure 2. Prevalence of T16189C polymorphism was found similar in patients (HTN, CTN, and PTC) and healthy control subject groups (13.8%, 16.1%, 20.8% and 17.3%, respectively) (*p*>0.05) (Table 1). Also, if the poly-C stretch prevalence was compared among the groups, no statistical difference was found (*p*=0.217). In regard to somatic mutation occurrence in np16189, 1 nodule (0.9%) of the 108 HTNs, 3 nodules (3.2%) in 95 CTNs and 7 nodules (14.58%) in 48 PTCs has somatic T16189C mutation (*p*=0.001) (Table 3). Moreover, excluding accompanied polymorphisms, which diminish the poly-C stretch, prevalence of the T16189C polymorphism alone is still significantly higher in PTC group the other groups (5/48, 10.4%) (*p*=0.002). Even, 4 of these 5 mutated tumoral tissues belong to micro-PTC group which may represents that could be a distinctive feature for the papillary thyroid micro-carcinomas rather than macro-carcinomas (Table 3).

### Comparison of all somatic mtMSIs together among the three study groups

The frequency of somatic alterations in D310 (39/251, 15.53%) was higher than in D514 (17/251, 6.7%), in D568 (4/251, 1.6%) and T16189C (11/251, 4.3%). Having a somatic

mutation at least one of the four mtMSIs was found to be higher in PTCs (17/48, 35.4%) than in HTNs (26/108, 24.1%) and CTNs (16/95, 16.8%) (*p*=0.046) (Table 4). Frequency of nodules harboring at least two somatic alterations was not statistically different among the groups studied (*p*=0.061). But in micro-PTC samples, this ratio was increased to 4/21 (19%) (*p*=0.012) (Table 4).

### Somatic mutations out of the mtMSIs in mtDNA control region

Totally 17 nodules in 251 (6.8%) samples have at least one somatic mutation out of mtMSIs described previously. In 5 HTNs, 9 somatic mutations and in 4 CTNs, 6 somatic mutations were detected whereas 19 somatic mutations detected in 8 PTCs. (*p*=0.010) (Table 3). If the HTNs and CTNs are compared with micro-PTCs and macro-PTCs separately, the prevalence of mutated nodules 3/27 (11.1%,) in macroPTCs was closed to benign nodules (*p*=0.338). In spite of that, the prevalence of mutated nodules 5/21 (23.8%) in microcarcinoma was the highest (*p*=0.002) (Table 3) (nodules were harbored with mutations in 14 different nucleotide position). All somatic mutations and frequencies were summarized in Supplementary Table 2.

## Discussion

Mutation and SNP prevalence in CR were investigated at HTN, CTN and PTC samples in this study. Prevalence of SNPs located at CR, tRNA^phe^, and 12sRNA genes in patients groups were compared with the healthy control subjects. However, the mtDNA haplogroups were determined in both patient and healthy control subject groups and the haplogroups and SNPs were evaluated for susceptibility/protective effects on thyroid entities.

Cancer is a multifactorial disease in which genetic and environmental factors play a role. On the other hand, mtDNA haplogroup variations have been shaped by environmental selection to adopt the environment. Thereby, naturally occurring mtDNA variations are not neutral. Moreover, the studies in literature reveal that the mtDNA variations were correlated with the metabolic and degenerative diseases. Also, they cause genetic susceptibility to the certain cancers. Thus, it confirms the functional importance of mtDNA variations (Wallace, 2016). However, Turkey is a bridge amongst Europe, Asia and Africa. Modern Turkish population has European H, U, J, T, K,W, I, V and X haplogroups, in addition to having Asian A, B, C, D, G, M, N and African L haplogroup (Guney et al., 2014; Mergen et al., 2004) as also confirmed by the present study. H haplogroup, which is a dominant haplogroup in Western-Euroasian populations, is also detected as the most common haplogroup (29.81%) in this study.

The number of studies related to the effects of mtDNA haplogroups on susceptibility to thyroid cancers is rather limited in the literature. However, in a study of 66 PTC cases in the Chinese population, Su et al., (2016) reported that A4, B4a and B4g mtDNA haplogroups were associated with PTC development. In spite of this, in a study conducted in 100 thyroid cancer cases, Fang H et al. (2010) reported that the D4a haplogroup had a higher risk of thyroid cancer in Chinese population. In addition, in a study of 114 thyroid follicular adenomas and 121 PTC cases, Cocos et al. (2017) found that K haplogroup have a protective effect on thyroid cancer in the Romanian population. In this study, the frequency of the K haplogroup in the healthy control group was found similar to that of Cocos et al., (2017) (6.73% vs 6.92%). On the contrary the frequency of the K haplogroup was found remarkably higher in thyroid patients (4.16%). It is close to the K haplogroup frequency of the healthy Turkish population. On the other hand, increased risk of benign and malign thyroid disease occurrence in U haplogroup was attracted attention in this study. Similar to our findings; Booker et al. (2006) also reported that U haplogroup has approximately 2-fold increased risk of prostate cancer and 2.5-fold increased risk of renal cancer in white North American individuals (Booker et al., 2006). Beside this, in patients with J haplogroup, the risk of developing benign thyroid nodules was found to be decreased. Also, in individuals with HV haplogroup, the risk of developing PTC was found to be lower.

In this study, 8 SNPs (T146C, G185A, C194T, C295T, G16129A, T16304C, A16343G and T16362C) located in the mtDNA CR were found to be associated with benign and malignant thyroid tumor occurrence. Also, it was observed that the distribution of 3 SNPs (C16296T C16355T and A16399G) within the patient groups was statistically different. All the SNPs were located in HVS1, HVS2 and HVS3. Besides, G16129A was located in the extended termination association sequence 1 (ETAS1) and T16304C was located in the ETAS2 which take place in mitochondrial replication (Sbisa et al., 1997). Amongst these SNPs, T16362C was found associated with PTC occurrence in Chinese population previously (Su et al., 2016). In this study, the prevalence of T16362C was detected higher in CTNs than HTN, PTC and NC (12.3% vs 1.5, 4.2 and 3.8%, respectively). If it is considered together with the findings of Su et al. (2016), in which the prevalence of T16362C was detected lower than the healthy controls (15.15 vs 30.00%) in the study of 66 PTC cases in the Chinese population, also, less than 1% of the HTNs and 5-15% of the CTNs have the potential to transform the malignant thyroid cancer, it might be suggest that presence of T16362C polymorphism has a protective effect on CTNs against malign transformation. However, it might be suggested that presence of the T146C and G185A polymorphisms indicate low risk of benign and malignant thyroid entities according to the obtained data. On the other hand, it might be stated that the presence of the C194C and G16129A SNPs causes susceptibility to PTC. In addition, C295T polymorphism was detected in all patients with J haplogroup, as it mentioned previously in literature this polymorphism is associated with the J group (Mueller et al., 2012; van Oven and Kayser, 2009). Also, it is found that T16304C and A16343G SNPs were associated with susceptibility to HTN formation. However, although C16296T polymorphism was detected in 9.2% of HTNs, it was not detected in PTC cases. Similar to that, C16355T and A16399G polymorphisms were not detected in the PTC cases but 8.9% of the individuals with CTNs were has one. Thus, it can be suggested that the presence of these polymorphisms has a protective effect against PTC, but large population based studies are needed to prove it. On the other hand Su et al., (2016) reported that G709A polymorphism on 12S RNA has a protective effect against PTC in a case control study. However, in this study, G709A polymorphism was found more frequently in the PTC cases than the healthy population, but statistically not significant (25% vs. 17.3%; *p*>0.05). Nevertheless, contrary to the findings of Su et al., (2016), it might be postulated that individuals with CTN carrying G709A polymorphism are more susceptible to have PTC in Turkish population, given the potential for carcinogenesis of cold thyroid nodules. But as already pointed out, large population based studies are required in different population to confirm it. Up to date, the biological significance and mechanism of action of these of 8 SNPs, which were found to be associated with PTC risk in this study, has not yet been clarified. However, these mtDNA polymorphisms may have contributed to the development of benign and malignant tumor phenotypes as a result of aerobic glycolysis induction with impaired ATP and ROS production which together with mitohaplogroups and environmental factors as previously reported in the literature (Zhou et al., 2007).

In this study, somatic mutations in the CR were identified in all benign and malign tumor samples studied. D310 was found to be hot spot for benign thyroid nodules consistent with the previous studies (Maximo et al, 2005; Ding et al., 2010). On the other hand, nps 16189 and 514 were detected as hot spots for PTC.

The first poly-cytosine repeat of the D310 region, a hotspot for primary tumors, is highly polymorphic and the distribution of the residues ranges from 7C to 9C in healthy populations (Aral et al., 2006; Chatterjee et al., 2011; Xu et al., 2012; Yacoubi-Loueslati et al., 2009). However, it is reported that D310 mutations are frequently occur in adenomas and premalign tumors in the literature consistent with the present study (Ding et al., 2010; Maximo et al., 2005; Xu et al., 2012). Therefore, it has been suggested that D310 mutations occur as a results of the combination of oxidative stress, low efficiency of the mtDNA mismatch repair mechanisms and weak proofreading ability of the polymerase γ (Mambo et al., 2003). For this reason, It has been claimed that D310 has an important role in maintaining the number of mtDNA copy number. The increase/decrease in the number of the C residues in the Poly-C tract may affect the rate of DNA replication by disrupting the binding of DNA polymerase and other trans-acting elements (Fliss et al., 2000). However, the functional contribution of D310 alterations in cancer development is still unclear. It is concluded that the different rate of D310 variations in the different tumor types suggest the existence of alternative mechanisms for the generation of some D310 alterations, such as the rate of acquired mutations during the tumor development and the number of mitochondria per cell. So that most D310 alterations limited in polymorphic range reveal that most D310 variants in tumors are unlikely to functionally impair mitochondria (Chatterjee et al., 2011).

However, on the other hand in a case-control study of breast cancer, it is reported that C6, C7, C10 and C11 repeat frequencies were higher in malignant tumor tissues contrary to the common poly C repeat (C8 and C9) in healthy Canadian population (Xu et al., 2012). Moreover, in the same study it is detected that the healthy surrounding tissues close to the tumor tissues, which are harboring with D310 somatic mutations had an identical genetically pattern whereas more than 2 cm away healthy surrounding tissue had C8 poly-C repeat in D310, which is common in the general population. The similar findings were observed in head and neck tumors and biliary cancers (Ha et al., 2002; Tang et al., 2004). Because of the fact that, although the authors are not of opinion that the mutations in the D310 are not related to cancer progression, starting out definition of “field cancerization" and considering mutations are early events in carcinogenesis; they postulated that alterations in D310 could be used as potential clonal expansions marker in the identification of genetic anomalies in premalign breast cancer cells.

In this study, C7 repeat at D310 was found higher in CTNs, micro-PTCs, macro-PTCs, and *BRAF*V600E (+) PTCs, whilst it was detected less frequent in the HTN than the general Turkish population (Figure 1). In addition, 3 *BRAF*V600E(+) accompanying with lymph node metastasis had C7 repeat at D310. One of the *BRAF*V600E(-) tumor samples accompanying with vascular invasion also had 1 base deletion as a somatic mutation (C8/C9→C7/C8/C9) at D310. Furthermore, most of the somatic mutations at D310 in the HTNs (75%) were 1/2 bp insertions. In spite of this, 1/2 bp deletions were more frequent in both CTNs (50%) and PTCs (80%). Thus, C7 repeat at D310 may provide a selective advantage for clonal proliferation to nodular tissues in terms of tumor progression in concordance with (Xu et al., 2012). Therefore, it might be suggested that the D310 alterations can be used as a potential clonal expansion marker in genetically altering cells in premalignant thyroid nodules.

Another mutational hot spot at the D-loop is T16189C. As it is mentioned previously, T16189C polymorphism has been associated with various multifactorial diseases and mtDNA copy number (Liou et al., 2010; Liu et al., 2003; Mueller et al., 2011; Tipirisetti et al., 2014). The wild type mtDNA sequence between np 16184-16193, is C_5_TC_4_. In a study of 837 healthy control subjects, Liou et al. reported that mtDNA copy number was lower in individuals with uninterrupted poly-C tract (>10 bp) between np 16184-16193 whereas mtDNA copy number was higher in individuals with various interrupted variants than the wild type individuals (Liou et al., 2010). However, in the literature it is postulated that the mitochondrial single strand binding protein (mtSSB) binds more efficiently to the interrupted poly-C tract than the uninterrupted one, and also, this sequence is located at 7S DNA binding sequence which plays pivotal role in the regulation of mtDNA synthesis. Thus, the uninterrupted variant may impair the mtDNA replication and reduce the mtDNA copy number (Tipirisetti et al., 2014). Additionally, it is suggested that reduced mtDNA content may affects the efficiency of the mitochondrial electron transport system (ETS), reduce the ATP/ADP ratio and increase ROS production (Mueller et al., 2011). However, increased ROS production may damage many cellular components including mtDNA. The damaged mtDNA, if not repaired properly, produces mtDNA mutations. These mutations may eventually lead to the initiation of tumorigenesis and sustain cancer development (Lu et al., 2009).

In this study, germline T16189C polymorphism were found to be similar in patients and healthy control subjects. The finding at healthy control group was also consisted with our previous study (Aral et al., 2011). However, there was no any statistical significance between the uninterrupted poly-C tract and benign and malign thyroid entity occurrences. Therefore, it can be concluded that neither the uninterrupted poly-C tract nor T16189C polymorphism causes the susceptibility to benign and malignant thyroid entities in Turkish population. But, interestingly, as previously mentioned in the results section and Table 3, somatic mutations at np 16189 were detected higher in PTC samples than the benign thyroid ones. Moreover, somatic mutation frequency was much higher in micro-PTC samples whereas it was found similar to CTNs in macro-PTC samples. Also, when the presence of uninterrupted poly-C tract was compared amongst the groups as a somatic alteration it was more frequent in micro carcinomas. Furthermore, somatic mutations at non-mtMSI sequence in micro-PTC group were detected much higher than the other groups and decreased in macro-PTC group (Table 3).

Fliss et al. (2000) hypothesized that somatic mutations may be lost during subsequent tumor oxygenation by replicative segregation with the cell turning back towards the more oxidative mtDNA genotype favoured in the metastatic environment. To support this hypothesis, Brandon et al. (2006) suggested that somatic mutations might be expected to arise and lost from the tumor cells at different times during the neoplastic process. The severely deleterious tumorigenic mutations that inhibit the ETS would be advantageous in the initial phases of tumor growth when the tumor requires mitochondrial H_2_O_2_ to drive cell proliferation. In this early stage, the tumor is hypoxic and thus may tolerate OXPHOS deficiency. However, when the tumor becomes vascularized and/or metastasizes and cells return to a high oxygen tension environment, then it may be more advantageous for the established transformed cells to revert a more oxidative metabolism (Brandon et al., 2006). Due to our study limited to the control region, we could not speculate that somatic mutations at control region are correlated with defects in ETS. But, we can suggest that such an adaptive somatic mutations, such as T16189C, occurring at D-loop may reflect the consequence of deleterious mutations in the coding region. Therefore, we can conclude that in this study, presumably, increased frequency of somatic T16189C mutation in micro-PTCs might be associated with reduced mtDNA copy number. In these micro carcinoma samples, T16189C mutation might induce internal ROS production by means of reducing mtDNA content. Thus it may provide selective adaptation to aerobic glycolysis in these tumor cells that have not yet either vascularized or metastasized and this would provide selective advantage to the tumor cells to initiate malignant tumor transformation in concordance with the literature.

However, the issue of mtDNA copy number in thyroid cancers is contentious. In a few studies it was demonstrated that mtDNA copy number is increased in thyroid cancers (Mambo et al., 2005; Su et al., 2016). But Reznik et al. (2016). indicated that there is no clear increase or decrease in mtDNA copy number when compared with normal thyroid tissues Although, T16189C is associated with reduced mtDNA copy number (Liou et al., 2010), none of these previous studies on thyroid cancer evaluated the effect of T16189C variant or uninterrupted poly-C tract on mtDNA copy number and a clear genotype-phenotype correlation is missing. Also, Mambo et al. (2003) suggested that alteration in mtDNA content might be an early event in thyroid carcinoma formation. But none of the studies in the literature compared the mtDNA content between thyroid micro-carcinomas and macro-carcinomas up to date. The findings on somatic T16189C variation and uninterrupted poly-C stretch in the present study indicates that further studies are necessary to determine the mtDNA copy number in micro and macro PTC samples harbored with T16189C variation or not in order to elucidate the role of this variant in thyroid tumor progression. In a further study we plan to compate mtDNA copy number at micro- and macro-carcinomas in PTC cell subtypes.

Another mutational hot spot on the mtDNA CR is D514, although the function of this CA repeat is not clearly understood. Variation rate in the CA repeat at D514 were detected in low frequency at endometrial, over, gastric cancers and gliomas, whereas it is high in breast cancers, head and neck cancers and thyroid cancers in literature (Liu et al., 2003; Maximo et al., 2005; Pang et al., 2008; Sharma et al., 2005). However, Kleist et al (2017) reported that genomic instability in D310, 514, 16184 in colorectal cancers is associated with lymph node metastasis rather than the primary tumor. Maximo et al. (2005) was found 33.2% of malignant tumors (29.4% of follicular carcinomas and 36.7% of PTCs) had a deletion/insertion at D514 while only 10% of adenomas had a dinucleotide alteration at D514 in a case-control study with 66 patients. Based on these findings, they suggested that the dinucleotide alterations in the D514 may be associated with tumor progression in thyroid cancers. In this study, it is detected that the somatic mutation rate in PTCs at D514 CA repeat was higher with respect to the other groups consistent with Maximo et al., (2005) (Table 3). In addition, as mentioned before the frequency of the wild-type 5CA repeat at D514 was found 85.6% in healthy control subjects and the frequencies of 4CA, 6CA, 7CA and 8CA repeats at D514 were found 7.7%, 2.9%, 2.9% and 0.9%. But the frequency of 4CA repeat increased to 37.3% in micro-PTC samples harboring with the somatic mutations whereas the surrounding tissue in these tumor samples has 13.2% 4CA repeat frequency similar to other benign tumor samples and healthy control subjects. On the other hand, the frequency of 4CA repeat was detected in macro-PTC samples (35.3%) similar to the micro-PTC samples whereas the 4CA frequency the surrounding tissue in these tumor samples increased to 24.1% remarkably different from the other groups (Figure 2) (Table 3). Therefore, the obtained data in this study confirms the findings of Maximo et al., (2005) and indicates that D514 mtMSI region might be used as a prognostic marker for the PTCs.

Wallace. theorized that the risk of cancer increases with the age and all age-related diseases such as degenerative diseases, cancer and aging shares a common underlying mitochondrial pathophysiology (Wallace, 2005). In the literature mitochondrial mutations and mtDNA instability associated with several human cancers as mentioned before. On the other hand, Schon et al., (2012) claimed that healthy mitochondria is crucial for tumor progression for adequate *de novo* pyrimidine synthesis. Therefore, this hypothesis brings forward the speculations that there is a relationship between mitochondrial genomic integrity and tumor aggressiveness or not, and the mitochondrial genome is a “genetic sanctuary” during the oncogenic process or not. This is the dilemma which is not properly answered in any research in literature. In thyroid carcinogenesis, PTC frequently develops from CTNs and micro- carcinomas may be considered as the initial phase of tumorigenesis. At the later stages of the disease, the tumor proceeds to macro-carcinoma and eventually metastasis and vascular invasion. *BRAF* V600E mutation is associated with poor prognosis, lymph node metastasis, recurrence and tumor aggressiveness of the PTC in the literature (Xing, 2013). In this study, prevalance of somatic mutation was detected higher in *BRAF*V600E(-) micro-carcinomas (57%) than the CTNs (17.9%), *BRAF*V600E(-) macro-carcinomas (30%) and *BRAF*V600E(+) cases (20%). Furthermore, no mtDNA somatic mutation was detected in metastatic *BRAF*V600E(+) tumors (n=3). It may speculate that in the early phases of thyroid tumor development dysfunctional mitochondria (due to mtDNA mutations and/or reduced copy number) may promote tumor formation. On the other hand, healthy mitochondria is necessary for later stages as previously mentioned. Therefore, the data obtained in this study on mtDNA CR region indicates that, further case-control studies with larger populations are required in different cell subtype PTC cases with different tumor stages to determine the contribution of mtDNA genome stability/instability and mtDNA copy number to the PTC tumorigenesis.

As a result, in this study it is found that the individuals having mtDNA U haplogroup are more susceptible to benign and malign thyroid entities and 8 mtSNPs are associated with benign or malign thyroid tumors in Turkish population. However, it has been detected that the D310 poly-C sequence may be a potential diagnostic marker for clonal expansion of premalignant thyroid nodules. However, when the previous study of Maximo et al., (2005) was taken together with this study, it was demonstrated that mtDNA D514 CA alterations can be used as a prognostic marker in thyroid cancers. In addition, this study provides clues that mitochondrial genomic alterations may play a critical role in the nodular transformation at different stages of papillary thyroid carcinogenesis. The further larger population based studies on entire mitochondrial genome and mtDNA copy number in benign and malign thyroid tumors are needed to confirm these findings.

## Acknowledgements

This work was supported by two grants of the Research Fund of the Tekirdağ Namık Kemal University, Project numbers NKUBAP.00.10.AR.14.07 and NKUBAP.00.10.AR.15.05.

